# Age-related remodeling of the glycocalyx drives T cell exhaustion

**DOI:** 10.1101/2024.12.06.627213

**Authors:** Hanlin Zhang, C Kimberly Tsui, Jesse Garcia Castillo, Audrey Evangelista, Esther Jeong Yoon Kim, Larry K Joe, Nicholas Twells, Ellen A Robey, Lara K Mahal, Michel DuPage, Andrew Dillin

## Abstract

Cell surface glycans, termed the glycocalyx, are essential regulators of cellular signaling and thus cellular development and functions, but how aging impacts the glycocalyx remains poorly understood. Here, using immune cells as a model system for studying the relationship between aging and glycocalyx remodeling, we show that α2,6-linked sialic acid – a terminal glycan epitope typically associated with inhibitory signaling – becomes downregulated in T cells from older animals. This downregulation is tightly correlated with age-associated accumulation of effector T cells, which are decorated with little to no α2,6-linked sialic acids. T cell aging renders older individuals more vulnerable to infections and cancers. To understand the role of α2,6-linked sialic acids in T cell physiology, we generated a mouse model with T cell-specific deletion of the sialyltransferase gene *St6gal1*. The chronic depletion of α2,6-linked sialic acids led to naïve T (T_N_) cells expansion in the periphery and premature T cell exhaustion. As a result, these mice were less able to control acute *Listeria* infection and chronic tumor growth. Blockade of the PD-1 pathway can partially restore the ability of *St6gal1*-deficient T cells to control tumor growth. Together, these data suggest that α2,6-linked sialic acids are critical for maintaining long-term T cell responsiveness, and the loss of α2,6-linked sialic acids may directly contribute to age-related T cell exhaustion.

## Introduction

Aging is accompanied by a decline of the immune system, termed immunosenescence^1^. This decline is linked with increased susceptibility to infections, autoimmune disease, and cancer. T cells are particularly susceptible to age-related dysfunction due to a combination of factors including thymic involution and clonal expansion in the periphery^2,3^. Without high levels of T_N_ production in the thymus, the total number of T cells and the proportion of naïve T cells (T_N_) available for responding to new pathogens declines significantly^4^. Increased conversion of T_N_ into memory (T_mem_) or effector (T_eff_) cells with age also occurs due to increased homeostatic proliferation and/or chronic antigen exposure^5,6^. Persistent exposure to activating signals may further result in T cell exhaustion, leading to upregulation of inhibitory receptors, such as PD-1, LAG-3, CTLA4, and TIM3, dampening T cell response against chronic infections and tumor cells^7^. Investigating mechanisms of cellular aging from distinct perspectives can help comprehensively understand T cell aging for restoring immune function and controlling age-related diseases.

The surfaces of all cells and organisms are decorated with a diverse repertoire of glycans, collectively known as the glycocalyx^8^. These glycans play critical roles in mediating both cell-autonomous and non-autonomous signaling events through interaction with membrane-bound and secreted glycan-binding proteins^9,10^. As immune cell development and activation are tightly controlled by extracellular signals, glycosylation can significantly impact immune cell functions. In T cells, glycans regulate thymic development, differentiation, activation, proliferation, and exhaustion^11–16^. Furthermore, specific changes in asparagine(N-)linked glycosylation on T cells have been shown to impair T cell function in aged individuals, in which the degree of N-glycan branching – modifications that negatively impact receptor clustering and activation – becomes upregulated on T cells of aged female mice and human^17^. How different types of glycosylation become altered with age and their functional consequences is starting to be unveiled.

Here, we investigated the changes in glycan on multiple immune cell types across age. We identify a previously unreported reduction in α2,6-linked sialic acid – a terminal glycan epitope typically associated with inhibitory signaling^18^ – specifically on aged T cells. We find that this reduction in α2,6-linked sialic acid is closely associated with activated T cells, which account for a larger proportion of total T cells with age. Using a conditional knockout mouse model that specifically depletes α2,6-linked sialic acid modifications in T cells, we discovered that chronic depletion of α2,6-linked sialic acid in T cells leads to T_N_ expansion in the periphery and premature T cell exhaustion. Interestingly, these results are in contrast to the acute removal of all sialic acids from T cell surfaces, which leads to more robust T cell activation and rejuvenates PD1^+^ exhausted T cells^19–22^, underscoring the importance of understanding the functional differences between sialic acid linkages as well as acute versus chronic alterations to T cell glycocalyx. Together, our results highlight the potential link between age-related loss of α2,6-linked sialic acid and T-cell exhaustion in immunosenescence and uncover the critical role of α2,6-linked sialic acid in mediating T_N_ maintenance and exhaustion.

## Results

### Profiling of glycocalyx changes during immunosenescence

To understand how aging affects the glycocalyx of immune cells, we measured the levels of different glycan motifs on monocytes, neutrophils, B cells, and T cells in the periphery blood from young (under 6 months) and old (over 18 months) mice (**See supplementary Fig. 1a for gating strategy**). Specifically, we assayed immature high mannose N-glycans, branched N-glycans, and sialic acids using lectins HHL, PHA-L, and SNA^23^ by flow cytometry (see table 1 for lectin binding specificities). This revealed a slight reduction in high mannose in B cells (**Supplementary Fig. 1b**), a reduction in complex branched N-glycans in neutrophils (**Supplementary Fig. 1c**), and most drastically, a loss of α2,6-linked sialic acid in both CD4+ and CD8+ T cells (**Fig. 1a and b**). As α2,6-linked sialic acid motifs are typically the capping sugar residue added to a LacNAc motif, (**Fig. 1c**), we measured the levels of unmasked LacNAc using ECL and found a corresponding upregulation in aged T cells (**Fig. 1d**). In addition, we also find an increase in α2,3-linked sialic acid using SLBR-N and SLBR-H (**Fig. 1e and Supplementary Fig. 1d**). However, different from ECL staining results where all SNA^low^ T cells are ECL+, the increase in α2,3-linked sialic acid was not found in all SNA^low^ cells (**Supplementary Fig. 1e and f**). These results indicate that the exposure of terminal LacNAc may directly result from reduced α2,6-linked sialic acid modification, but age-related upregulation of α2,3-linked sialic acid modification may have a distinct mechanism.

**Figure 1.**
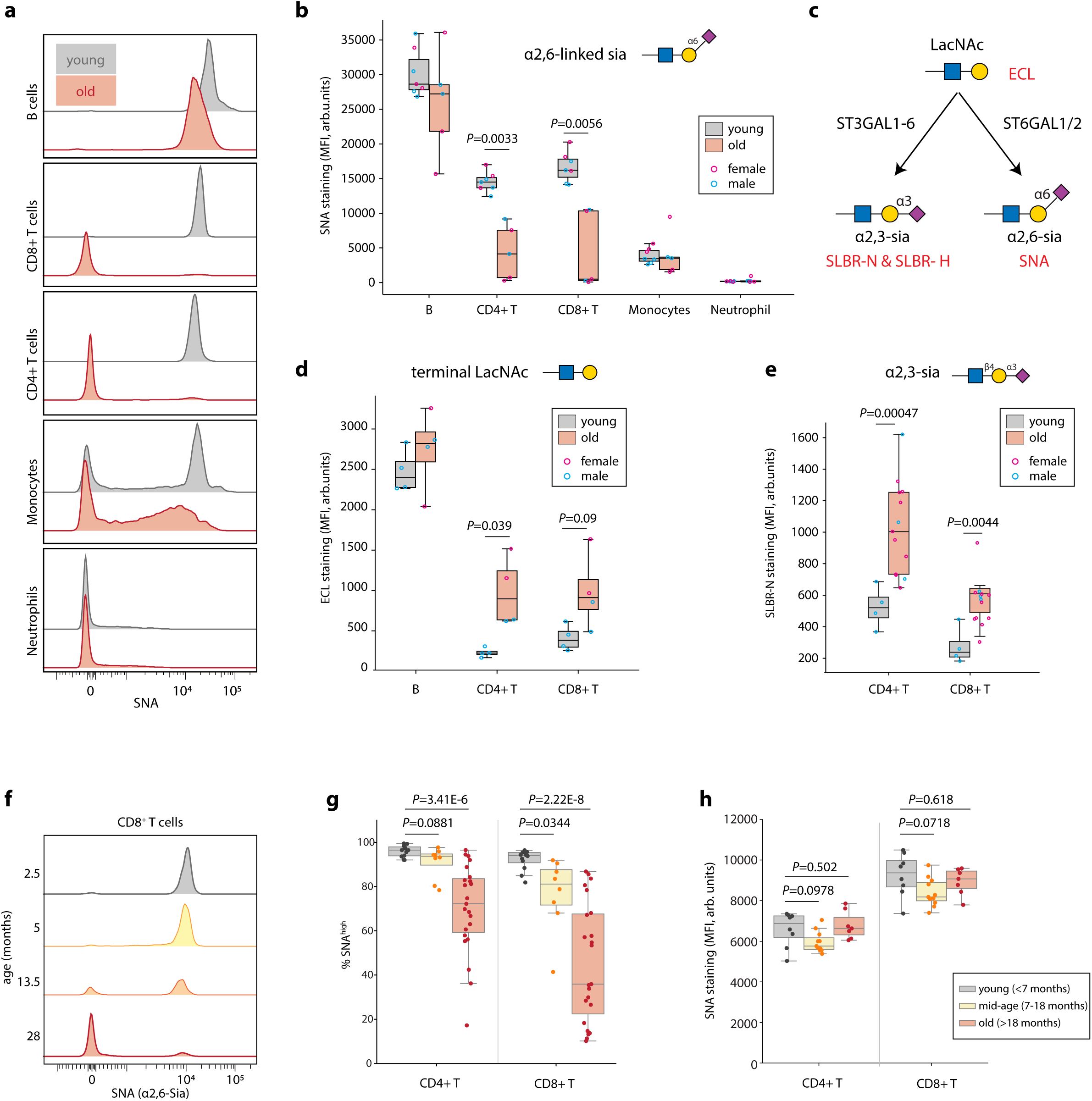
Aging is associated with reduced α2,6-linked sialic acid modification on T cells. **a/b.** Peripheral blood was collected from young (< 7 months, n=7) and old (>18 months, n=5) mice followed by SNA flow cytometry staining for measuring α2,6-linked sialic acid modifications. B cells (B220+), CD4+ T cells (CD4+), CD8+ T cells (CD8+), monocytes (Ly6C+), and neutrophils (Ly6G+) were gated based on their respective surface markers. One representative plot is shown in (a) (young = 1.5 months, old = 25 months) and summarized in (b). **c.** Biosynthetic pathways for sialic acid linkages. Lectin specificities are indicated in red. **d.** Peripheral blood samples from young (n = 4) and old (n = 4) mice were analyzed by ECL using flow cytometry staining. **e.** Peripheral blood samples from young (n=4) and old (n=13) mice were analyzed by SLBR-N using flow cytometry staining. **f-h.** Peripheral blood was collected from young (< 7 months, n = 12), mid-age (7-18 months, n = 8), and old (>18 months, n = 23) mice followed by SNA flow cytometry staining. One representative plot of CD8+ T cells staining is shown in f. The levels of α2,6-linked sialic acid were quantified by percent of SNA^high^ cells. **f-h.** Peripheral blood was collected from young (< 7 months, n = 8), mid-age (7-18 months, n = 11), and old (>18 months, n = 7) mice followed by SNA flow cytometry staining. The mean fluorescence intensity (MFI) of SNA^high^ cells were quantified. All data are represented in box plots showing the quartiles of the data with whiskers showing the rest of the distribution. *P* values were calculated by two-tailed student’s t-test

To gain better time resolution of this sialic acid change, we assayed α2,6-linked sialic acid across age – from 2 to 27 months old across immune cell types (**Supplementary Fig.1g**). Interestingly, instead of a gradual linear decrease of SNA staining on T cells, we find that as mice age, they gain a distinct population of T cells without SNA staining (**Fig. 1f**). Therefore, we decided to categorize T cells into SNA^high^ or SNA^low^ populations and analyze them separately. We find that as mice pass the middle age of 18 months, the population of SNA^low^ T cells starts to expand quickly with an increased variability among the aged population, mirroring the increased phenotypical and functional variability among T cells within the same-aged individual (**Fig. 1g**). This is especially true in CD8+ T cells, in which the vast majority of CD8+ T cells become SNA^low^ only after 18 months old. Interestingly, the levels of α2,6-linked sialic acid remain unchanged in the SNA^high^ T cells across age (**Fig. 1h**), further highlighting that the reduction of α2,6-linked in T cells is driven by the loss of this SNA^high^ population.

### Loss of α2,6-linked sialic acid with age is associated with T cell activation

Given that T cell activation has been shown to downregulate α2,6-linked sialic acid^14^, we hypothesized that the age-related loss of SNA^high^ CD8+ T cells is related to increased activation of T cells. We first tested whether activation of young T cells will result in the loss of α2,6-linked sialic acid. In concordance with previous reports, *ex vivo* activation by anti-CD3/anti-CD28 and *in vivo* activation by *Listeria monocytogenes* infection both result in the reduction of the proportion of SNA^high^ and a corresponding appearance of SNA^low^ CD8^+^ T cells (**Fig. 2a, b, and c**), reminiscent of changes we see in aged mice.

**Figure 2.**
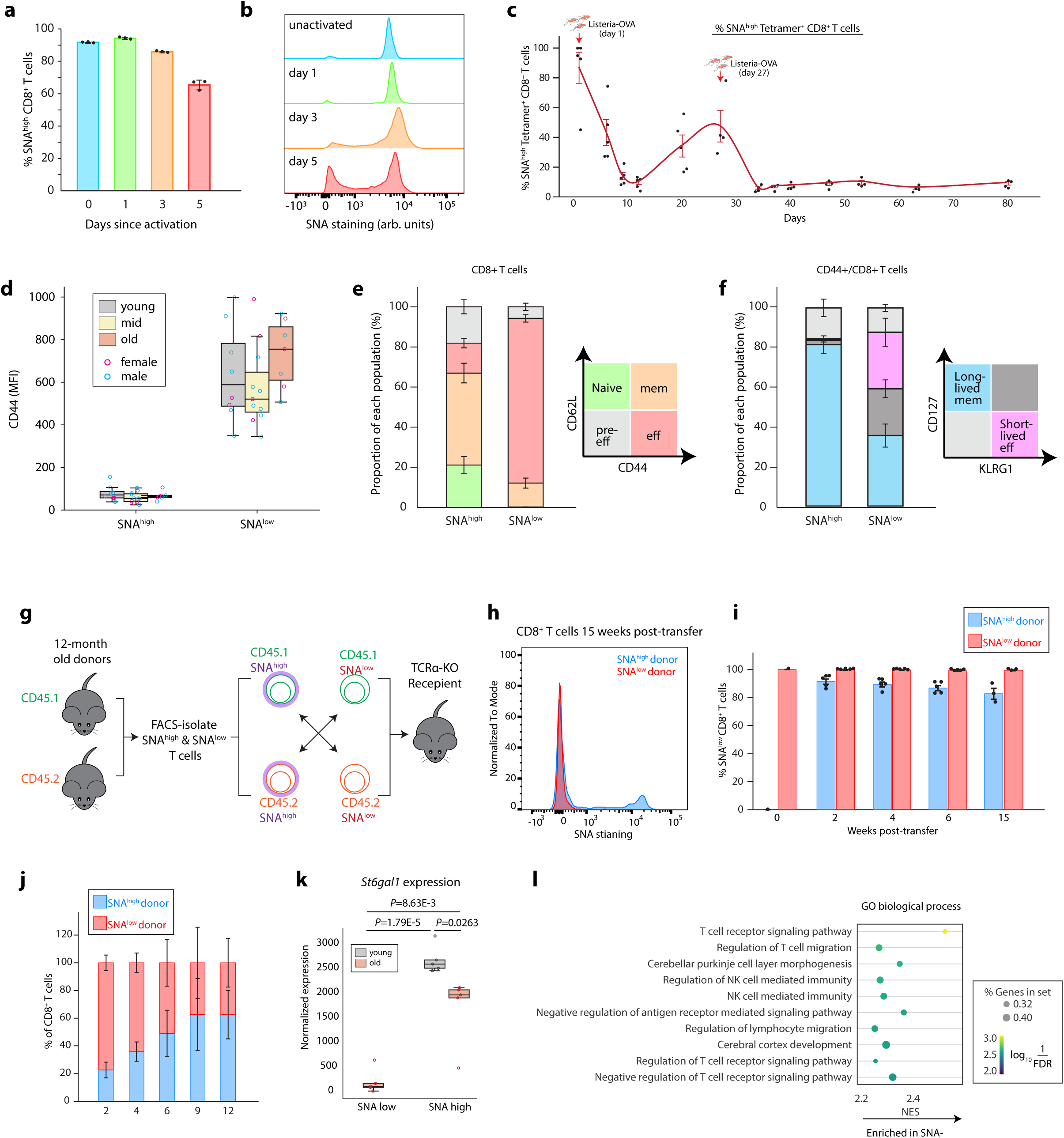
Loss of α2,6-linked sialic acid with age is associated with T cell activation. **a/b.** Splenic T cells from young mice were stimulated with anti-CD3/anti-CD28 in a 96-well plate for indicated days for SNA flow cytometry staining. Statistical summary (a) and a representative plot (b) is shown. n = 3. **c.** Young wildtype mice were infected with Listeria-OVA through *i.v.* injection on day 0 and day 28 respectively. Peripheral blood samples were collected on indicated days for measuring SNA of OVA tetramer-specific CD8^+^ T cells. n = 5. **d.** The levels of CD44 were compared between SNA^high^ and SNA^low^ CD8+ T cells from young (n=8). Mid-age(n=11) or old (n=7) mice. **e.** SNA^high^ and SNA^low^ CD8+ T cells from mid-age mice were analyzed based on CD44 and CD62L staining to assess naïve, pre-effector (pre-eff), effector (effector memory, eff), and memory (central memory, mem) subpopulations. n = 10. **f.** SNA^high^ and SNA^low^ CD44+ CD8+ T cells from mid-age old mice were analyzed based on CD127 and KLRG1 staining to assess short-lived effector (KLRG1+CD127-) and long-lived memory (KLRG1-CD127+) subpopulations. n = 10. **g-j.** SNA^high^ and SNA^low^ CD8+ T cells were FACS-isolated from CD45.1+ or CD45.2+ mice respectively, mixed at 1:1 ratio, and adoptively transferred into TCRα-KO recipient mice (g). Peripheral blood samples were collected at indicated time points for SNA staining. A representative plot is shown in (h) and quantified results are shown in (i). The competitive contribution to the reconstituted population from different donors is quantified in (j). n = 5 **k/l**. SNA^high^ and SNA^low^ T cells were sorted from young (only having SNA^low^ cells) and old mice for deep RNA-seq analyses. The expression of the *St6gal1* gene is shown in (k) and the GO pathway analysis result is shown in (l). n = 5. All data represented as mean ± SEM. *P* values were calculated by two-tailed student’s t-test

To directly test whether the age-related sialic acid changes may result from activation, we co-stained T cells from young and old mice with a panel of activation markers. We find that regardless of age, SNA^low^ CD8^+^ T cells are positive for the activation marker CD44 (**Fig. 2d**). Moreover, within the CD44^+^ antigen-experienced T cell population, SNA^low^ cells are mostly CD44^+^/CD62L^-^ effector T cells, while SNA^high^ cells are more likely to be CD44^+^/CD62L^+^ memory T cells (**Fig. 2e**). To determine whether these SNA^low^ CD8+ T cells are long-lived effector memory or short-lived effector cells, we additional stained CD127 and KLRG1 and found that a significant ratio of SNA^low^ CD8^+^ T cells are short-lived effector T cells (CD127^-^/KLRG1^+^), whereas the SNA^high^ CD8^+^ T cells are primarily long-lived memory or memory-precursor cells (CD127^+^/ KLRG1^-^)^24^ (**Fig. 2f**).

To more directly evaluate the developmental relationship between SNA^high^ and SNA^low^ T cells, *i.e.* whether they can transform to each other, we performed a reconstitution experiment by mixing SNA^high^ T cell from CD45.1^+^ mice and SNA^low^ T cells from CD45.2^+^ mice at the 1:1 ratio and adoptively transferred them into a TCRα-KO recipient mice that lack endogenous T cells (**Fig. 2g**). We find that during 15 weeks of reconstitution, SNA^low^ T cells were unable to regain α2,6-linked sialic acid, whereas SNA^high^ T cells can efficiently give rise to SNA^low^ T cells within 2 weeks of reconstitution while retaining a small proportion of SNA^high^ T cells (**Fig. 2h and i**). In line with our CD127/KLRG1 staining results, we found that the SNA^low^ CD8^+^ T cells proliferated faster than their SNA^high^ counterpart in recipient mice, but were relatively short-lived (**Fig. 2j**). Therefore, consistent with surface marker co-staining results, this functional adoptive transfer experiment further indicates that SNA^high^ T cells are stem cell-like naïve/memory T cells, whereas SNA^low^ T cells are terminally differentiated effector T cells.

As the α2,6-linked sialic acid modification recognized by SNA is mainly generated by the sialyltransferase ST6GAL1, we re-analyzed publicly available single-cell RNA sequencing data of young and old murine immune cells^25^ to determine whether we can detect any differences in *S6t6gal1* expression in young and old T cells (**Supplementary Fig. 2a and b**). In concordance with our SNA staining results (**Fig. 1a and b**), *St6gal1*-expression is enriched in B cells and, to a lesser degree, T cells, but not in monocytes or neutrophils. However, we were unable to detect sufficient *St6gal1* expression in the single-cell data even in the naïve T cell population that should all have high levels of α2,6-linked sialic acid, possibly due to the sequencing depth limitation. Therefore, to further understand the differences between the SNA^high^ and SNA^low^ populations of T cells, we performed bulk RNA-seq to compare these populations of T cells in young and old mice. We find that SNA^high^ T cells have higher expression of *St6gal1* relative to SNA^low^ cells, regardless of age (**Fig.2k**), indicating that the loss of α2,6-linked sialic acids is due to transcriptional downregulation of *St6gal1*. When comparing SNA^high^ and SNA^low^ T cells in aged mice, Gene-set enrichment analysis further revealed signatures of increased T cell receptor signaling and proliferation in SNA^low^ T cells (**Fig. 2l, Supplementary Fig. 2c and d**). Together, these results suggest that the accumulation of activated T cells with age results in the loss of the SNA^high^ T cell population, with reduced *St6gal1* gene expression as the direct cause at the molecular level.

### Loss of α2,6-linked sialic acid leads to excessive expansion of mature T cells in periphery

To study the developmental and functional consequences of the loss of α2,6-linked sialic acid in T cells, we generated a T cell-specific *St6gal1-KO* mice to mimic activation- and age-related down regulation of *St6gal1* by crossing the *St6gal1^fl/fl^*mice with *CD4-Cre* transgenic mice^26^. As expected, T cells in these *St6gal1-KO* mice have specific depletion of α2,6-linked sialic acid only in their T cells, but not other immune cell types (**Supplementary Fig. 3a**). They also have increased ECL staining, which binds to the unmasked LacNAc structure that underlies α2,6-linked sialic acids (**Supplementary Fig. 3b**). Interestingly, α2,3-linked sialic acid levels are upregulated in *St6gal1-KO* CD4^+^ but not CD8^+^ T cells, suggesting that there might be a cell type-specific compensation effect during the sialylation process (**Supplementary Fig. 3c**).

Previous works have reported that T cell-specific depletion of all sialic acids by knocking out the sialic acid synthesis gene *Cmas* led to unaltered thymic T cell development but severe lymphopenia in the periphery with significantly reduced number of mature T cells^27^. In contrast, whole body knockout of *St6gal1*, which depletes the α2,6-sialic acids modification alone but in a non-tissue specific manner, led to reduced thymic cellularity and reduced DN, DP, and SP thymocytes^28^, indicating that proper sialic acid modification of most likely niche stromal cells is required to maintain thymic T cell survival. Complimentary to these systems, T cell-specific *St6gal1-KO* can be used for properly mimicking age-related reduction of α2,6-linked sialic acid on T cells and testing how sialylation may directly regulate T cell behaviors in an autonomous manner. Interestingly, deletion of *St6gal1* at the DP stage does not have a significant impact on post-DP stage thymic T cell development, as the proportions of DP and SP CD4^+^ and CD8^+^ thymocytes as well as their subpopulations all remain unchanged (**Fig. 3a**, **Supplementary Fig. 3d-h**). Therefore, consistent with previous reports, our data indicate that sialylation on the T cell surface is dispensable for T cell development and survival during their development in the thymus.

**Figure 3.**
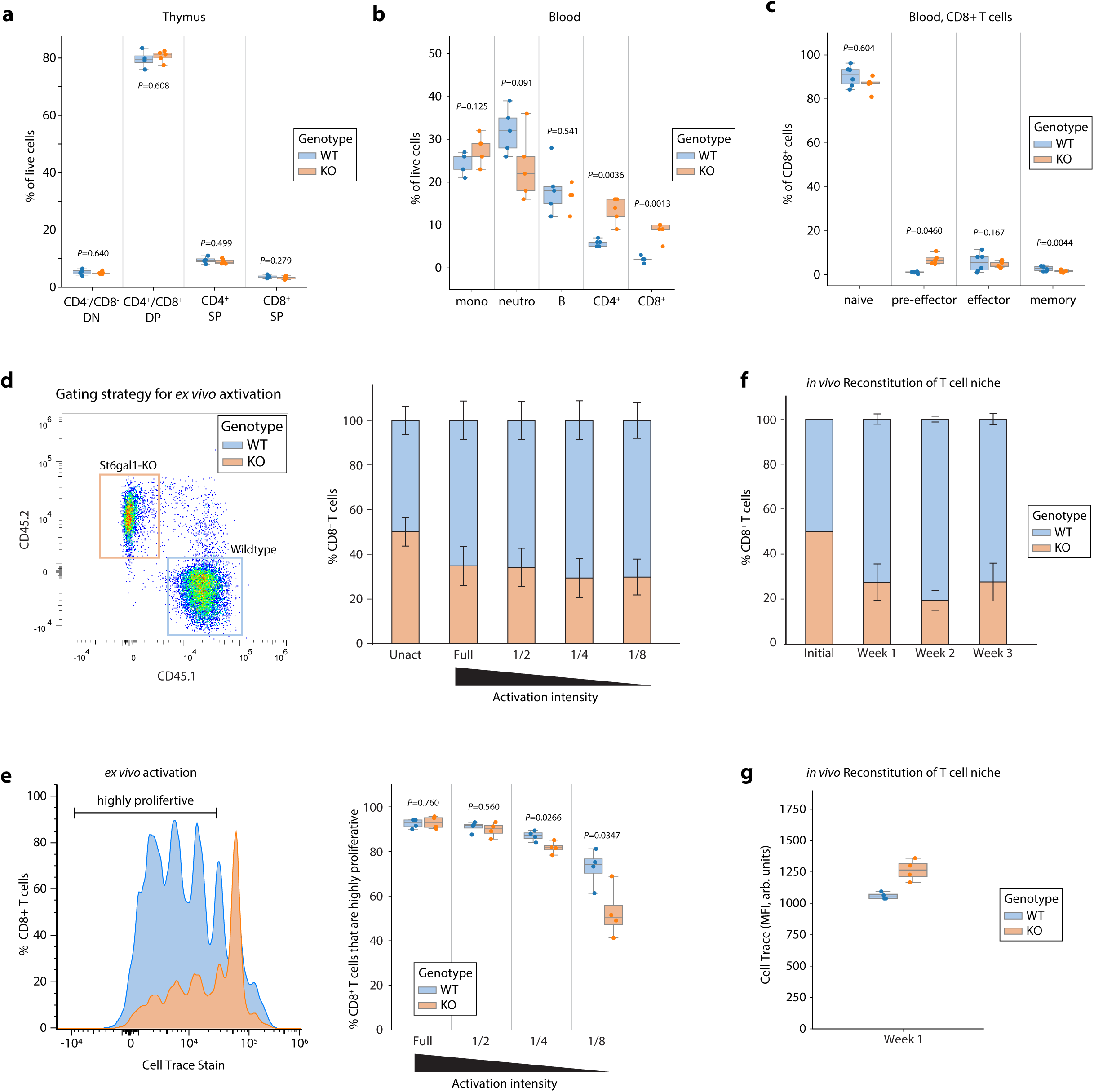
Loss of α2,6-linked sialic acids leads to expansion of mature T cells that are less responsive. **a-c.** Thymocytes (a), peripheral blood (b), and blood CD8^+^ T cells (c) from CD4-Cre, St6gal1^fl/fl^ mice were analyzed using flow cytometry. n = 4-5. **d-e**. CD8^+^ T cells from wildtype (CD45.1^+^) or CD4-Cre, St6gal1^fl/fl^ (CD45.2^+^) mice were mixed at the ratio of 1:1 for *ex vivo* activation using full dosage of or diluted anti-CD3/anti-CD28 for 3 days (d). Cell proliferation was measured using CellTrace Violet staining with a representative plot shown in (e) and quantified in (f). n = 4. Data represented as mean ± SEM. **f-g.** CD8^+^ T cells from wildtype (CD45.1^+^) or CD4-Cre, St6gal1^fl/fl^ (CD45.2^+^) mice were mixed at the ratio of 1:1, stained with CellTrace Violet, and adoptively transferred to TCRα-KO recipient mice. Blood samples were collected at indicated timepoints to assess reconstitution efficiency (g) and proliferation (f). n = 4. Unless otherwise noted, all data are represented in box plots showing the quartiles of the data with whiskers showing the rest of the distribution. *P* values were calculated by two-tailed student’s t-test.

Interestingly, a significant increase of mature CD4^+^ and CD8^+^ T cells in the peripheral blood was observed (**Fig. 3b**), suggesting that loss of α2,6-linked sialic acids may drive the expansion of mature T cells. To test whether deprivation of α2,6-sialic acid is sufficient to activate T cells, we examined the relative proportion of naïve (CD62L^+^CD44^-^), effector (CD62L^-^CD44^+^), and memory (CD62L^+^CD44^+^) CD8^+^ T cells (**Fig. 3c**). No significant changes were observed among these three populations. However, an emergence of the rare population of CD62L^-^CD44^-^ T cells was observed (**Fig. 3c**), which may mark a transient status of pre-effector T cells during early T cell activation^29^), indicating that SNA^low^ T cells may have received more stimulating signals which predisposes them but is not sufficient to drive their full activation. Consistently, T cell-specific *St6gal1-KO* does not lead to increased overall inflammation through measuring serum cytokines using an ELISA array (**Supplementary Fig. 3i**). Overall, T cell-specific loss of α2,6-linked sialic acid leads to expansion of mature CD4^+^ and CD8^+^ T cells possibly due to increased sensitivity to tonic TCR signals and homeostatic cytokines. These expanded T cells remain phenotypically naïve.

### Loss of α2,6-linked sialic acid leads to premature T cell exhaustion

To more directly test the functional consequence of α2,6-linked sialic acid deprivation on CD8^+^ T cells, we first measured the ability of *St6gal1-KO* and wildtype CD8^+^ T cells to respond to anti-CD3/CD28 stimulation *ex vivo*. To do so, we stimulated *St6gal1-KO* (CD45.2^+^) CD8^+^ T cells with anti-CD3/anti-CD28. As a stringent control, WT (CD45.1^+^) cells were included in the mixture so that the stimulation for different genetic backgrounds can be identical within each well. Unexpectedly, *St6gal1-KO* T cells failed to respond as efficiently as wildtype T cells (**Fig. 3d**). Indeed, under weaker stimuli, *St6gal1-KO* T cells proliferated remarkably less than their wildtype counterpart (**Fig. 3e**). In line with the *ex vivo* activation results, *St6gal1-KO* T cells also were less proliferative and less efficient in reconstituting the empty niche in TCRα-KO mice *in vivo* (**Fig. 3f and g**).

Next, we assessed how the specific loss of α2,6-linked sialic acid impacts T cell function in various immune activation models *in vivo* (**Fig. 4a**). We first used *Listeria* as an acute infection model, which typically provides a very strong stimulus for T cell activation ^30^. In line with our *ex vivo* findings, *St6gal1-KO* T cells have greatly dampened primary response, generating less than half as many antigen-specific CD8+ T cells when compared to wildtype mice (**Fig. 4b and c**). The corresponding memory response is also weaker (**Fig. b and Supplementary Fig. 4a**). Together, these results suggest that the loss of α2,6-linked sialic acid causes T cells to be less responsive and produces a dampened immune response.

**Figure 4.**
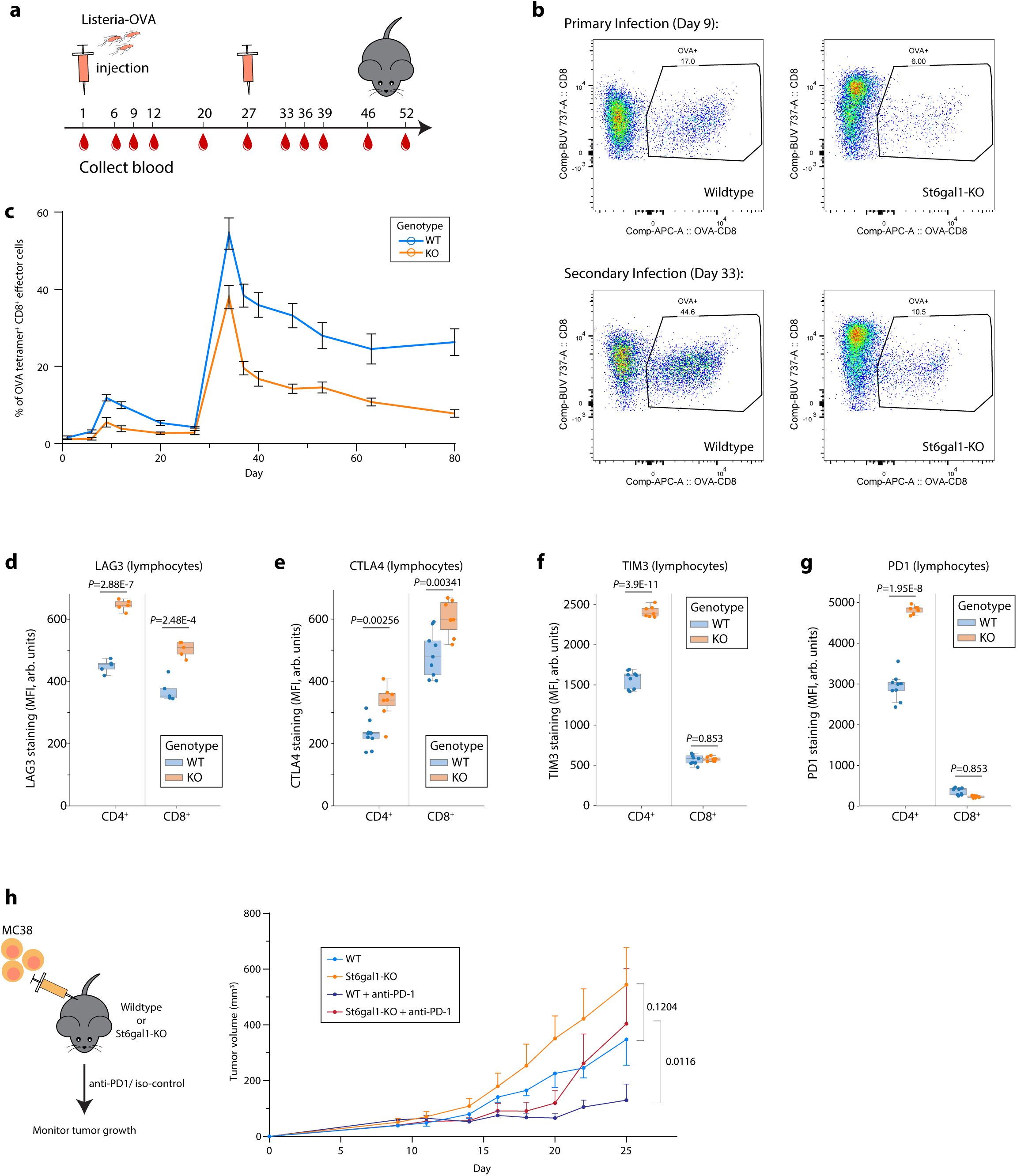
Loss of α2,6-linked sialic acids results in prematurely exhausted cells. **a-c.** Wildtype or CD4-Cre, St6gal1^fl/fl^ mice were infected with Listeria-OVA on day 0 and day 28 and blood samples collected on indicated days (a). OVA antigen-specific CD8^+^ T cells were analysed using flow cytometry (b) and quantified in (c). n = 5-6. Data represented as mean ± SEM. **d-g.** T cell exhaustion markers including LAG3 (d), CTLA4 €, TIM3 (f), and PD1 (g) were quantified on T cells in peripheral blood from wildtype or CD4-Cre, St6gal1^fl/fl^ mice. N = 5-9. Data are represented in box plots showing the quartiles of the data with whiskers showing the rest of the distribution. *P* values were calculated by two-tailed student’s t-test **a. h.** MC38 tumors were engrafted subcutaneously to wildtype or CD4-Cre, St6gal1^fl/fl^ mice followed by anti-PD1 treatment starting from day 11. Tumor volume was measured with time. N = 5-6. Data represented as mean ± SEM. Two-way ANOVA with post hoc Dunnett’s test.

Given that *St6gal1-KO* T cells are less responsive to activation stimuli both *ex vivo* and *in vivo*, we hypothesized that the loss of α2,6-linked sialic acid may cause T cells to become more exhausted due to increased sensitivity to persistent low-grade stimulation of tonic TCR signaling and homeostatic cytokines. Indeed, circulating *St6gal1-KO* T cells express increased exhaustion markers LAG3, CTLA4, TIM3, and PD1 (**Fig. 4d-g**). As T cell exhaustion is particularly relevant to chronic stimulation such as during tumor progression, we tested whether *St6gal1-KO* T cells were also less efficient in controlling tumor growth in an MC38 tumor engraftment model. Consistently, tumors in T cell-specific *St6gal1-KO* mice grow significantly faster than those in wildtype mice. Remarkably, anti-PD1 treatment efficiently suppressed tumor growth in these *St6gal1-KO* mice during the first week of treatment (**Fig. 4h**). This indicates that the loss of α2,6-linked sialic acids on T cells causes exhaustion that can be rescued by checkpoint inhibitor treatment in the short-term. However, after 9 days of treatment, anti-PD-1 lost its efficiency in *St6gal1-KO* mice, indicating that multiple exhaustion pathways may be involved in driving the exhaustion of T cells lacking α2,6-linked sialic acid.

## Discussion

Dysregulation of T cells is a key contributor to the declining immune system with age, causing older individuals to be more susceptible to a wide array of infectious, autoimmune, and malignant diseases ^2,3^. In this study, we investigated how the glycocalyx, which intimately regulates cellular development and function, changes with age on immune cells, particularly CD8^+^ T cells. Utilizing lectin staining on different immune cell types in young and old mice, we identified a dramatic reduction in α2,6-linked sialic acid modifications specifically in T cells from older animals. Additional experiments reveal that this reduction is linked to age-associated accumulation of T_eff_ cells, which are decorated with little to no α2,6-linked sialic acids. Particularly, these T cells cannot give rise to T cells with high α2,6-linked sialic acids, indicating that this is a lineage-restricted population possibly associated with terminal differentiation. Using a conditional T cell-specific *St6gal1-KO* mouse model, we further discovered that the loss of α2,6-linked sialic acid leads to excessive T cell expansion and exhaustion, causing T cells to be less responsive to activating stimuli. Mice without α2,6-linked sialic acid on their T cells are less able to control *Listeria* infection and tumor growth. Finally, we show that these exhausted T cells can be partially rescued by the blockade of the PD-1 axis, highlighting the importance of α2,6-linked sialic acids in preventing T cell exhaustion.

In contrast to our findings, previous works have shown that the acute removal of all sialic acid linkages by sialidases on T cells enhances T cell activation^19,21^. However, our data suggests that chronic depletion of α2,6-linked sialic acid can be detrimental to T cell activity and lead to premature exhaustion. We hypothesize that the long-term removal of α2,6-linked sialic acid sensitizes T_N_ cells to tonic TCR signaling and homeostatic cytokines, leading to excessive expansion and exhaustion of T cells. Future work investigating the differences between acute and chronic depletion of sialic acids on T cells will be important as many sialic acid-modifying therapeutics are being explored for cancer and CAR-T treatments^31–33^. Furthermore, α2,6-linked sialic acid modification may directly modify inhibitory receptors such as PD1, LAG3, TIM3, and CTLA4, which are all glycosylated. In particular, CTLA4 signaling and cell surface retention has been shown to be regulated by N-glycan branching^34^. How or whether these inhibitory receptors are regulated via α2,6-linked sialic acid modification is unknown and warrants further investigation.

Age-related reduction of α2,6-linked sialic acid modification was found in both murine CD4^+^ and CD8^+^ T cells. Our current research primarily focuses on CD8^+^ T cells, what about CD4^+^ T cells? It has been reported that antigens modified with sialic acids promote the generation of regulatory T (Treg) cells and suppress IFN-γ-producing inflammatory CD4^+^ T cells^35^, raising an intriguing possibility that sialylation may also impact CD4^+^ T cell signaling and functions. It begs further investigation how sialylation changes during CD4^+^ T cell differentiation and what the acute and chronic impacts of desialylation on different subpopulations of CD4^+^ T cells are. Importantly, dendritic cells become more tolerogenic upon taking sialylated antigens with reduced generation of inflammatory cytokines for priming CD4^+^ T cells^35^. Therefore, sialylation may modify T cell functions via both T cell intrinsic and extrinsic mechanisms. Interestingly, while human and murine T cells do not have exactly the same glycocalyx profiles or glycan remodeling dynamics^36^, the α2,6-linked sialic acid modification has been shown to also decline in all T cell subsets including CD8^+^, CD4^+^ and γδ T cells upon viral infections in humans^37^. Further comparison between different T cell subtypes in both human and murine models can generate valuable knowledge about how glycans guide sophisticated T cell dynamics and how evolution adopts appropriate strategies of immune cell glycan remodeling to fit different species.

Phenotypical biomarkers are useful tools for predicting T cell efficacy in vaccination and immune therapy. A variety of surface markers have been identified as an indicator of different status of T cells, such as CD44, KLRG1, PD1, and LAG3 ^38^. Indeed, some widely used T-cell markers function as receptors for glycans, including CD44^39^and CD62L^40^. This indicates that glycan-mediated signaling is intimately involved in regulating T cell homeostasis and activation. Our study revealed that α2,6-linked sialic acid depletes when T cells enter a terminally differentiated effector status, and that this depletion can drive T cell exhaustion. Therefore, SNA staining can be used as a novel binary marker for T cell activation, effector lineage restriction, and early exhaustion. We also noticed that age-related reduction of the SNA^high^ T cell ratio varies a lot among animals over 18 months old (Figure 1h). It is possible that the efficacy of their CD8^+^ T cells against infection and tumors also varies remarkably in this elder population, which can be predicated by α2,6-linked sialic acid levels as indicated by SNA staining.

In conclusion, our data demonstrate that α2,6-linked sialic acid depletion as indicated by SNA staining can be used as a novel marker for T cell aging in murine models. This change occurs during CD8^+^ T cell activation and is irreversible under physiological conditions. T cells with low α2,6-linked sialic acid are more prone to exhaustion and less efficient in controlling infection and tumors. Future investigations focused on how α2,6-linked sialic acid directly impacts T cell aging and exhaustion in human immune cells may provide additional biomarkers and therapeutic strategies for modulating T cell functions.

## METHODS

### ANIMAL MODELS

C57BL/6J wild-type mice (000664, JAX), CD45.1+ B6.SJL mice (002014, JAX), *Tcra-KO* mice (002116, JAX), *CD4-Cre* mice (022071, JAX), and *St6gal1-floxed* mice (006901, JAX) were purchased from the Jackson Laboratory. All mice were bred and maintained at the Berkeley Office of Laboratory Animal Care (OLAC) facility until they reach indicated ages for experimental use. Animal experiments were approved by the local ethical review committee and performed under project licenses AUP-2020-01-12908-1 and XXX.

### METHOD DETAILS

#### Flow Cytometry

Cells were stained with fixable aqua Live/Dead staining (L34966, Thermo Fisher), FcR block (Thermo Fisher), and surface marker antibodies in PBS containing 2 mM EDTA at 4 °C for 20 minutes and analyzed with a five-laser LSR Fortessa X-20. Specifically, fluorophore-conjugated lectins such as SNA (21500045-1, Bioworld, 1:200),and ECL (FL-1141-5, Vector Laboratories, 1:200), SLBR-N (made in-house in Mahal lab), were stained together with other surface marker antibodies. To stain OVA antigen-specific CD8^+^ T cells, the OVA (SIINFEKL)-MHC I tetramer (Cat, 1:500) and FcR block were diluted in PBS-EDTA and mixed with cells for staining at room temperature for 1 hour first, followed by staining of other surface markers at 4 °C. FOXP3 intracellular staining was performed using the FOXP3 Staining Buffer Set (00-5523-00, Thermo Fisher) following manufacture’s protocol. Acquired data were analyzed using FlowJo 10.8.

#### Adoptive Transfer of T cells

SNA^high^ or SNA^low^ CD8+ T cells were FACS sorted from mid-aged CD45.1^+^ or CD45.2^+^ donor mice and mixed at the 1:1 ratio. A total of 500,000 donor T cells were adoptively transferred into young *Tcra-KO* recipient mice by *i.v.* injection. Peripheral blood samples were collected from recipient mice at indicated timepoints. For transferring WT CD45.1^+^ and *St6gal1-KO* CD45.2^+^ T cells, splenocytes were collected from age-matched young donors by smashing and filtering the spleen tissue through a 70 µm cell strainer and red blood cell lysed. Splenocytes were then analyzed using FACS to measure CD8^+^ T cell ratios, and mixed to keep CD8^+^ T cells at the 1:1 ratio. A total of 10,000,000 donor splenocytes were adoptively transferred into young *Tcra-KO* recipient mice followed by peripheral blood sampling at indicated timepoints.

#### Mouse T Cell Stimulation

Cell culture plates (96 well, 3596, Corning) were coated with 1 µg/mL (full dose, or diluted to 1/2, 1/4, 1/8) of anti-CD3 antibody (16-0032-81, Thermo Fisher) in PBS at 4 °C overnight. On the following day, blood cells were collected from WT CD45.1^+^ and *St6gal1-KO* CD45.2^+^ mice, mixed at the ratio of 1.5:1 (WT: *St6gal1-KO*, so that the ratio of CD8^+^ T cells can be closer to 1:1), followed by red blood cell lysis and CellTrace staining. Mixed blood cells were then resuspended in the T cell culture medium that contains RPMI1640 (containing HEPES, 22400105, Thermo Fisher), 10% Fetal Bovine Serum (FBS), Penicillin/Streptomycin (15-140-122, Thermo Fisher), GlutaMAX (35050-061, Thermo Fisher), 50 µM β-mercaptoethanol (M3148, Sigma), and 3 µg/mL (full dose, or diluted to 1/2, 1/4, 1/8) anti-CD28 (MA110172, Thermo Fisher), and cultured in the pre-coated plate for 3 days. About 25 uL of blood was cultured in each well of a 96-well plate. Mixed cells were also analysed on day 0 to determine the original ratio of CD8^+^ T cells for normalization purpose later.

#### CellTrace staining

To measure cell proliferation, CellTrace Violet (C34557, Thermo Fisher) was used according to the manufacturer’s protocol. In short, cells were resuspended in 1 mL CellTrace Violet staining solution (5 µM working concentration, dissolved in PBS) in a 15 mL Falcon tube, and incubated in dark at room temperature for 20 minutes. Then the staining solution was topped up with RPMI washing media (RPMI 1640 with 10% FBS) and incubated for 5 minutes to quench the free dye. After centrifugation, cells were washed once more with RPMI washing media and resuspended in appropriate T cell culture medium that contains indicated amount of anti-CD28.

### Listeria Infection

#### Tumor Engraftment

##### RNA-seq analysis for SNA^high^ and SNA^low^ T cells

SNA^high^ and SNA^low^ CD8+ T cells from mouse thymus of five young and five old mice were isolated by FACS. RNA was purified using an RNeasy Mini Kit (74106, Qiagen) and stored at −80°C until further processing. RNA-Seq library preparation and NovaSeq PE150 sequencing were performed by MedGenome. RNA-Seq analysis was performed by uploading.fastq files to the Galaxy web platform, using the public server at usegalaxy.org. and mouse FASTA reference transcriptome GRCm39 was used for alignment. Tools included Kallisto Quant v0.46.2+galaxy and DESeq2.

##### Reanalysis of single-cell RNA sequencing results

Metadata file and UMI counts were obtained directly from YV Teo et al., 2023 ^25^, and analyzed using ScanPy^41^. Only cells expressing 200-4000 genes with less than 10% of mitochondrial reads were included in subsequent analysis. Default parameters of ScanPy were used unless otherwise specified. UMAP and Leiden clustering were performed using ScanPy (n_neighbors=10, n_pcs=30, and resolution = 0.5). The same cell markers as YV Teo et al. 2022 were used for identifying cell types. They were Cd3e for all T cells, with Cd4 for CD4+ T cell and Cd8a for CD8+ T cells, Cd79a and Ms4a2 for B cells, Nkg7 for NK cells, Ly6c2 and Cx3cr1 for monocytes or dendritic cells (DC), C1qa, C1qb and C1qc for macrophage, Fcer1a and Cd200r3 for basophil and Hba-a1 for erythrocytes. The expression levels of St6gal1 were plotted on the same UMAP.

#### Cytokine Array

Serum cytokines were measured using the Proteome Profiler Array (Mouse Cytokine Array Panel A, ARY006, BioTechne) according to the manufacturer’s protocol. In short, the array membranes were blocked at room temperature for 1 hour. In the meantime, serum samples collected from mice of indicated genotypes were diluted in the Array Buffer at the ratio of 200 µL : 1.3 mL, mixed with 15 µL of reconstituted Mouse Cytokine Array Panel A Detection Antibody Cocktail, and incubated at room temperature for 1 hour. After blocking, the sample/antibody mixture was added to the array membrane and incubated at 4 °C overnight on a rocking platform shaker. After washing, the array membrane was incubated with the IRDye 800CW Streptavidin secondary antibody (926-32230, LiCor) and detected using the LiCor Odyssey imaging system. Signals were quantified using Fiji.

### Statistics and Reproducibility

All data are presented as mean + s.e.m. or with box and whisker plots showing quartiles and distributions. Experiments are representative of at least two independent experiments unless otherwise indicated. Two groups were compared using the two-tailed Student’s *t* test in Excel. Two-way ANOVA with post hoc Dunnett’s test was used for comparing tumor volumes and calculated in Prism. No data were excluded from analyses.

---

**Figure S1.**
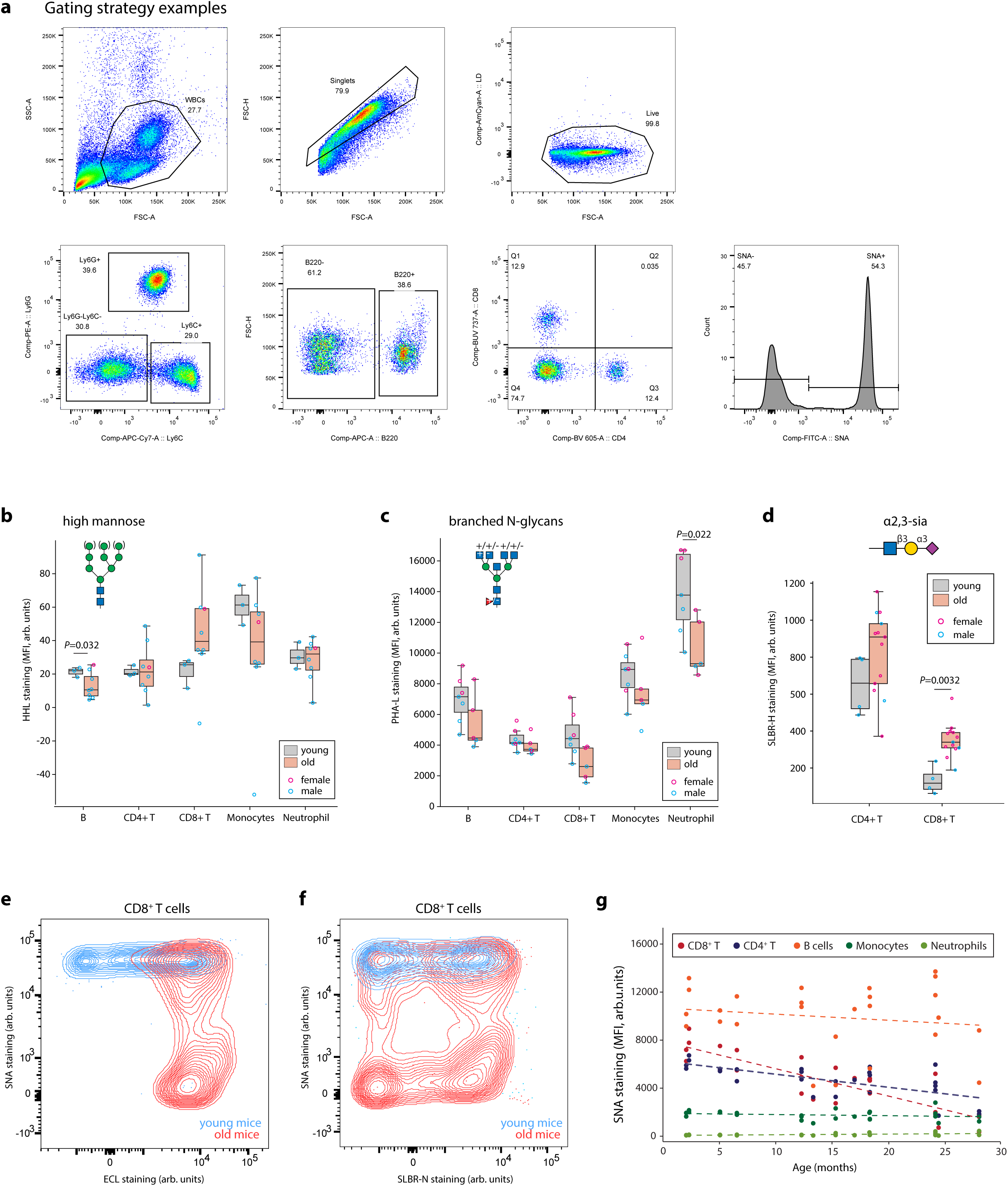
Glycan profiling of immune cells during aging. **a.** Representative flow cytometry graphs of gating strategy for different immune cell types **b-d.** Peripheral blood was collected from young (< 7 months) and old (>18 months) mice followed by HHL (b), PHA-L (c), and SLBR-H (d) flow cytometry staining for measuring high mannose, branched N-glycans, and α2,3-linked sialic acid modification respectively. n =3-8 **e-f.** A representative plot of SNA/SLBR-N (e) or SNA/ECL (f) co-staining of CD8^+^ T cells from young and old mice. **g.** Peripheral blood samples from mice of indicated ages were analyzed for α2,6-linked sialic acid modification by SNA flow cytometry staining. n = 7-11. Data are represented in box plots showing the quartiles of the data with whiskers showing the rest of the distribution. *P* values were calculated by two-tailed student’s t-test.

**Figure S2.**
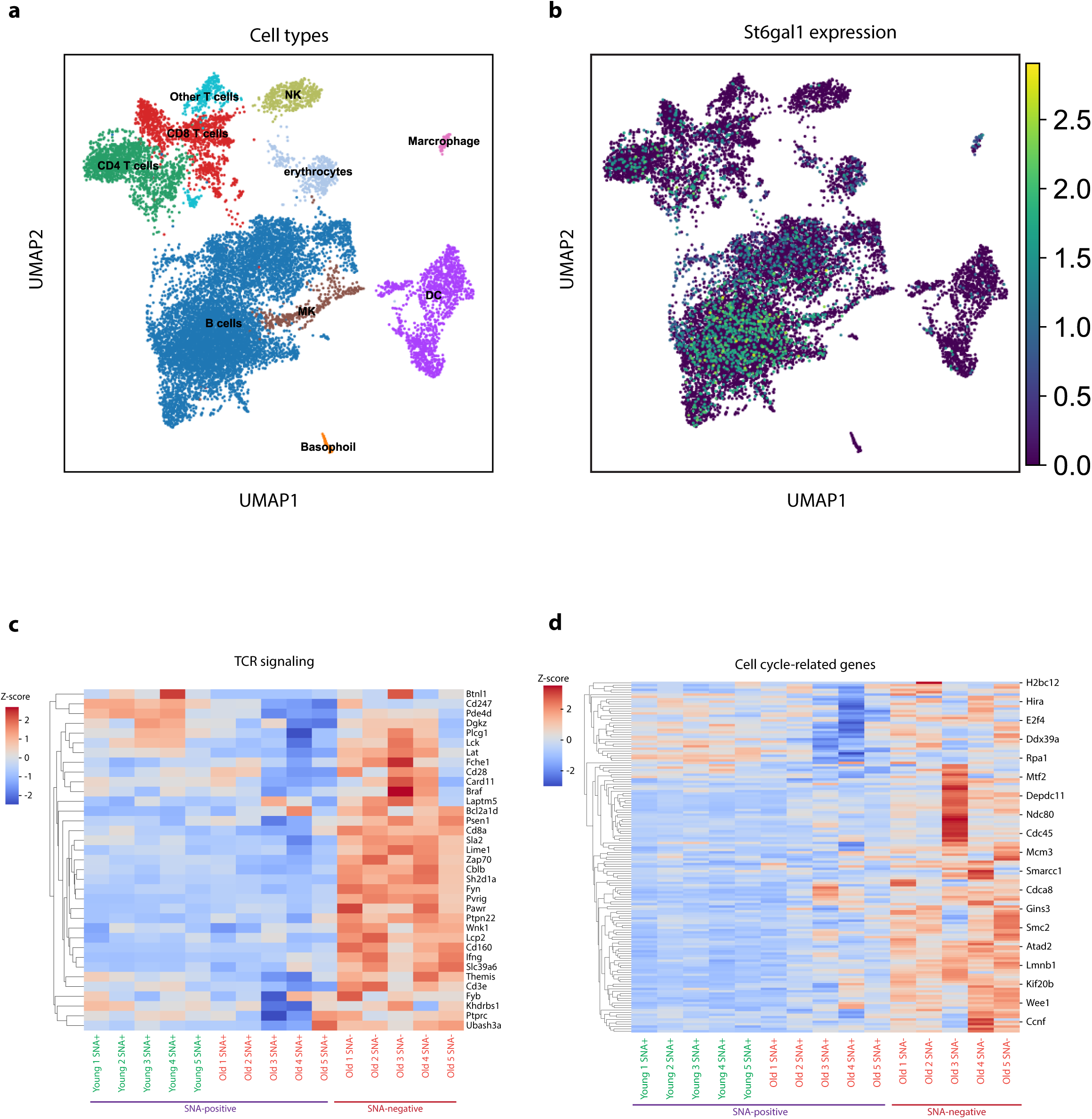
Subpopulation analysis of young and old immune cells using single cell RNA-seq and bulk RNA-seq. **a.** Single-cell RNA-seq data of immune cells from young and old mice blood were analyzed using UMAP. **b.** The expression of *St6gal1* was highlighted in the UAMP. **c/d.** SNA^+^ and SNA^-^ T cells (including both CD8+ and CD4+) from young and old mice were FACS-sorted for bulk RNA-seq analysis. Most significantly changed pathways are shown, such as TCR signaling (c) and cell cycle-related genes (d). n =5.

**Figure S3.**
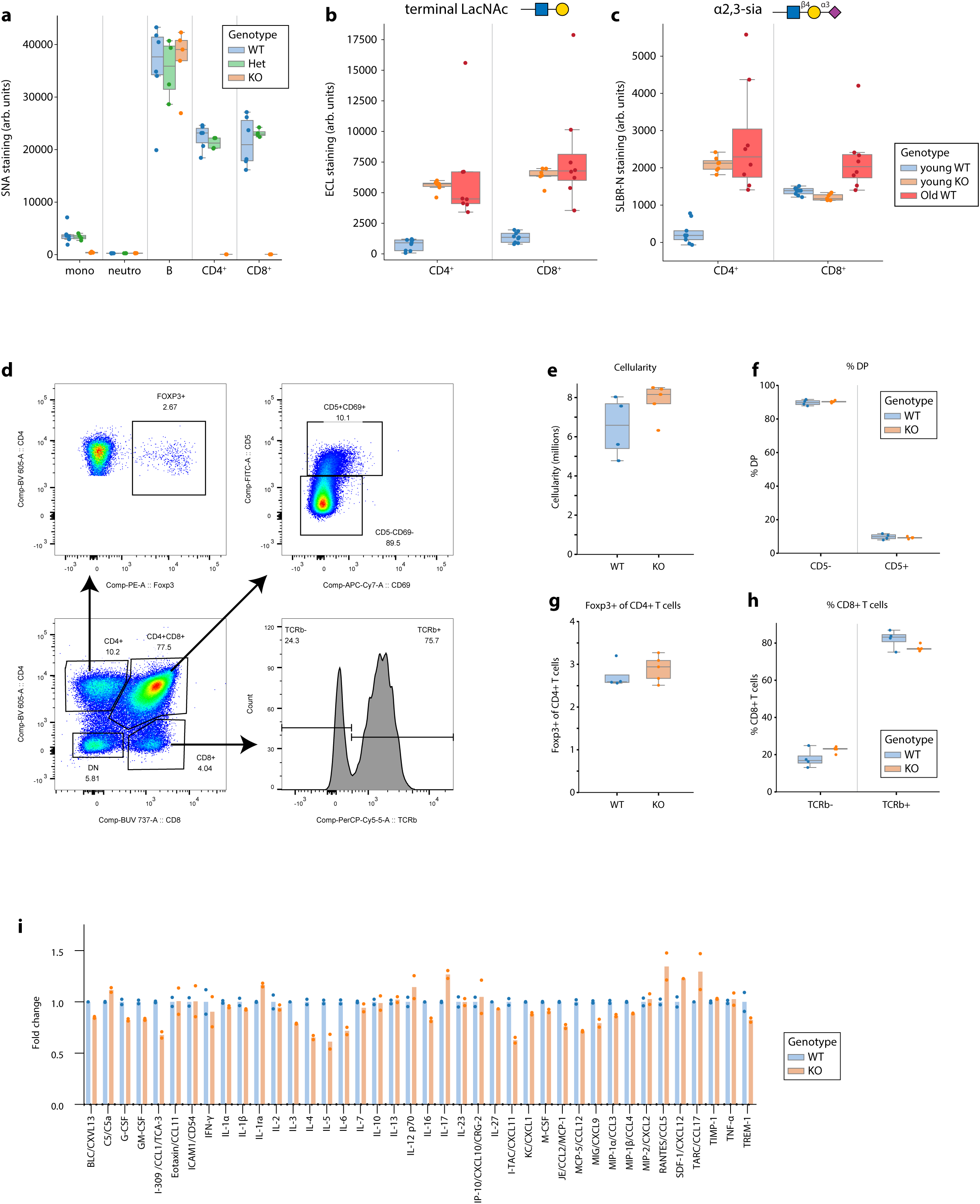
T-cell specific knockout of *St6gal1* does not significantly affect T cell development. **a.** Peripheral blood samples were collected from wildtype (WT, littermate control), *CD4-Cre, St6gal1-flox/wt* (Het), or *CD4-Cre, St6gal1-flox/flox* (KO) mice and stained with SNA for flow cytometry analysis. n = 4-6 **b/c.** Peripheral blood samples were collected from young WT, young KO, or old WT mice and stained with ECL (b) or SLBR-N (c) for flow cytometry analysis. n = 7-9 **d-h.** Thymocytes from WT or KO mice were analysed using flow cytometry. An example of gating strategy is shown in (d). Total cellularity (e), early (CD5^-^) and advanced (CD5^+^) double positive (DP) T progenitors (f), formation of Foxp3^+^ regulatory T cells (g), and immature (TCRb^-^) or mature (TCRb^+^) single positive (SP) CD8^+^ T cells (h) were quantified respectively. n = 4-5. **i.** Serum samples from young WT or KO mice were collected for a cytokine array assay. n = 2 (technical replicates). Data are represented in box plots showing the quartiles of the data with whiskers showing the rest of the distribution. *P* values were calculated by two-tailed student’s t-test.

**Figure S4.**
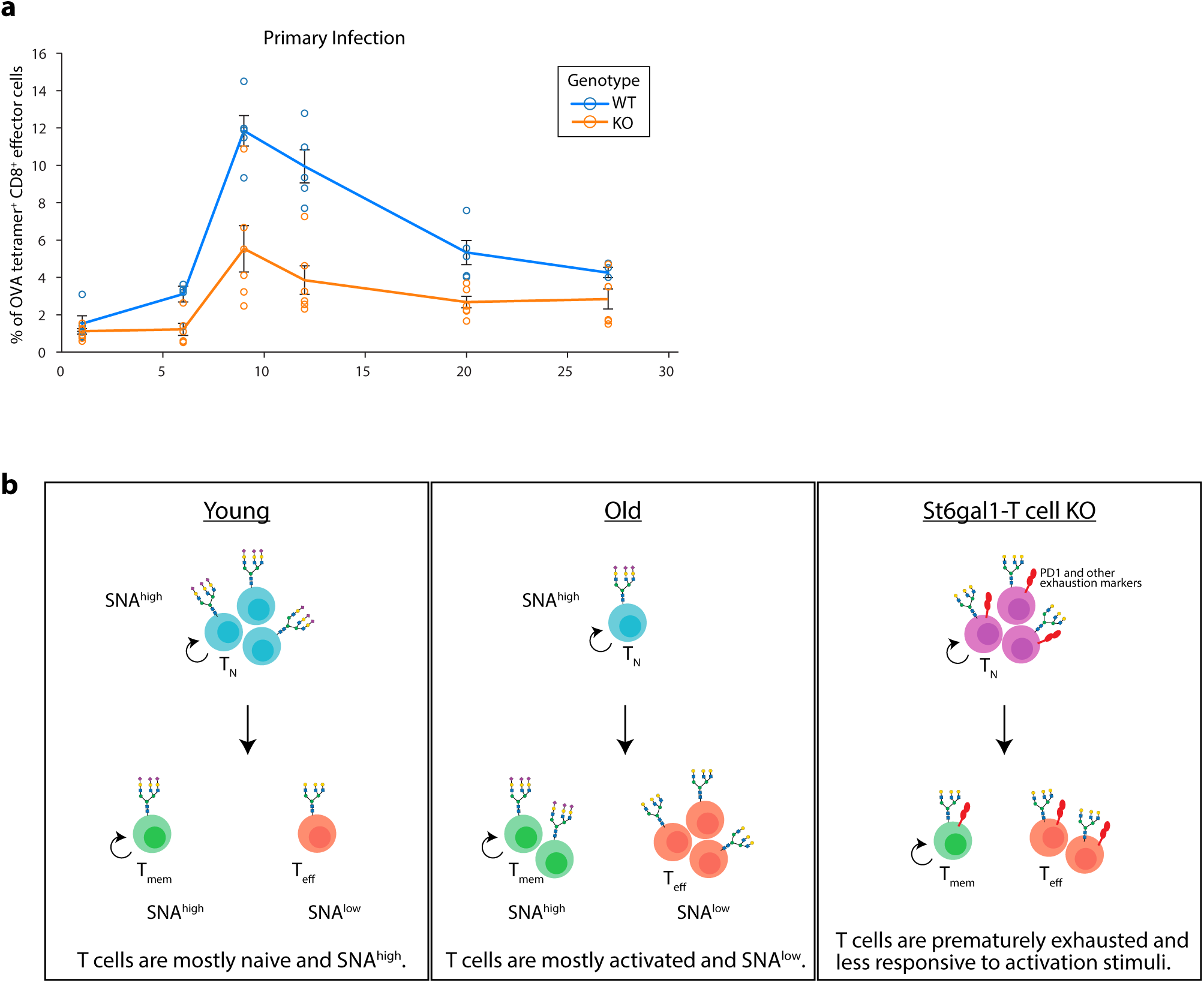
Loss of α2,6-linked sialic acids results in compromised effector CD8^+^ T cell responses. **a.** WT or *CD4-Cre, St6gal1^fl/fl^* (KO) mice were infected with Listeria-OVA on day 0 and blood samples collected on indicated days. OVA antigen-specific CD8^+^ T cells were analysed using flow cytometry. This panel is derived from Figure 4c and focuses on the primary response stage. n =5-6. **b.** A proposed working model of T cell aging-associated loss of α2,6-linked sialic acid. In young individuals, the majority of CD8^+^ T cells are naïve T cells with high levels of α2,6-linked sialic acid coating. In old individuals, the majority of CD8^+^ T cells become terminally differentiated effector T cells that are associated with depletion of α2,6-linked sialic acid. This loss of α2,6-linked sialic acid is not merely a consequence of T cell activation, but also directly affects T cell functions by driving their exhaustion.

